# Macrophage immunosenescence prolongs intraocular inflammation in aged mice via impaired induction of regulatory T cells

**DOI:** 10.64898/2026.01.15.697975

**Authors:** Taku Yamamoto, Keitaro Hase, Joseph B. Lin, Shinobu Yamaguchi, Norihiko Misawa, Jiayi Li, Kenta Kato, Ryo Terao, Brian S. Sohn, Daniel Du, Mitsukuni Yoshida, Charles W. Pfeifer, Tae Jun Lee, Jason Colasanti, Andrea Santeford, James T. Walsh, Rajendra S. Apte

## Abstract

Immune-mediated intraocular inflammation, called uveitis, is a leading cause of global blindness, with the highest burden of visual impairment falling on older individuals. Immunosenescence, the functional changes in immune cells with aging, impacts the age-associated immune response, but how immunosenescence and the molecular regulators of the age-associated immune response affect the clinical course of uveitis remains unclear. In the murine model of experimental autoimmune uveitis (EAU), aged mice demonstrated a delayed onset and peak of intraocular inflammation compared to young mice. In contrast to the canonical monophasic inflammation that rapidly resolves in young mice, aged mice developed persistent, chronic inflammation. Transcriptomic and flow-cytometric analyses of immune cells and the receptor-ligand interactome revealed a dominant macrophage-CD4^+^ T cell signature. This signaling pathway was functionally altered on both ends: macrophages from aged mice had an impaired capacity to generate peripherally induced regulatory T cells (pTreg) through an IL-6 regulated pathway, while CD4^+^ T cells co-cultured with aged macrophages demonstrated increased proliferation. Our study establishes aging as a key regulator of the effector immune response in uveitis. Regulatory T cells, specifically pTreg, are essential for resolving inflammation in uveitis and an impaired ability to induce pTreg led to a sustained, chronic inflammatory uveitis phenotype in old mice, thereby linking immunosenescence to persistent neuroinflammation. These findings highlight potential therapeutic avenues for vision-threatening uveitis, especially in older patients.

## 1 Introduction

Visual transduction begins when light enters the eye and reaches the neurosensory retina through a clear visual axis where it is perceived by photoreceptor neurons. This signal transduction cascade that begins in the retina culminates in the visual cortex and leads to a crisp image of the outside world. Inflammation inside the eye can affect this entire pathway by impairing light penetration to the retina, phototransduction within the retina, and proper transmission of information to the visual cortex. To protect the delicate neurovascular unit and preserve visual acuity, the eye has evolved as an immune-privileged environment where exuberant immune responses are actively suppressed (Wu et al., 2024). When these mechanisms are breached, the resulting intraocular inflammation, known as uveitis, is a devastating disease that accounts for about 10–15% of blindness in the US (Rothova et al., 1996). In immune-mediated, non-infectious uveitis, the effector immune response can cause unbridled inflammation (Maghsoudlou et al., 2025; Paley et al., 2024; Yang et al., 2022). Control of uveitic inflammation through immune regulatory mechanisms or therapeutic agents is crucial in order to prevent tissue damage and irreversible vision loss.

Of interest, visual prognosis in patients with uveitis varies with age. Although the prevalence of uveitis in older patients is not higher than in younger patients, older patients with uveitis are significantly more likely to experience vision loss compared to younger patients (Maini et al., 2004), but the etiology of why older individuals bear a disproportionate burden of vision loss from uveitis is unclear. Immunosenescence, the functional changes in immune cells with aging, is characterized by quantitative and qualitative changes in innate and adaptive immune cells, and it significantly impacts the age-associated immune response (Liu et al., 2023). This phenotype is also referred to as inflammaging (Franceschi et al., 2018). While adaptive immunity has long been a focus of immunosenescence, we have recently demonstrated a key role for macrophage senescence and innate immunity in the molecular pathogenesis of age-related macular degeneration (AMD), a classic aging-associated ocular disease (Apte et al., 2006; Kelly et al., 2007; Terao et al., 2024). Numerous studies have also revealed altered immune effector functions in cellular chemotaxis and phagocytosis associated with aging (Moss et al., 2023). We were interested in examining whether impaired immune regulation in age-associated uveitis was responsible for persistent inflammation and vision loss and elucidating the cellular components of the immune system that contributed to this age-associated phenotype.

In order to further explore inflammaging in uveitis, we utilized the well-described murine experimental autoimmune uveoretinitis (EAU) model of non-infectious uveitis that represents a T cell-mediated immune response to a retina-specific antigen, interphotoreceptor retinoid-binding protein administered by subcutaneous injection (Caspi et al., 2008). The key immune cell mediators are thought to be CD4^+^ T cells and myeloid cells. In the EAU model, young mice develop an acute monophasic inflammatory response that peaks around two weeks and resolves within five weeks (Agarwal et al., 2012). Here, we uncovered an interesting delay in the initial inflammation peak in old mice. Furthermore, the inflammatory response in old mice was biphasic, with a delayed initial peak and a persistent, low-grade inflammation following the initial peak that did not resolve throughout the observation period. Mechanistically, transcriptomic and flow-based analyses demonstrated that a macrophage-CD4^+^ T cell interaction dominated the infiltrating immune landscape. During EAU, old mice demonstrated significantly higher levels of CD4^+^ T cell proliferation while macrophages from old mice had a significantly reduced capacity to generate regulatory T cells (Tregs), resulting in an EAU response in old mice that was biphasic and chronic. These findings support the concept that macrophage immunosenescence contributes to aberrant CD4^+^ T cell activity that leads to sustained, chronic intraocular inflammation and is a potential mechanism for increased vision loss associated with uveitis in older individuals.

## 2 Results

### 2.1 Aging leads to delayed onset but sustained intraocular inflammation

Longitudinal EAU clinical scores in young (7-week-old male, n = 6) and old mice (20-month-old male, n = 8) were assessed by ophthalmic biomicroscopic examinations from 10 to 68 days after immunization. The peak EAU clinical score in young mice was 2.5 ± 0.56 at 16 days after immunization. In contrast, old mice demonstrated a peak clinical score of 1.8 ± 0.20 at 29 days after immunization, thus demonstrating a delay in the peak of inflammation in old mice as well as a blunted peak amplitude of response compared to young mice (Figure 1a). Following these mice longitudinally, EAU scores were acute and monophasic in young mice as described previously (Kerr et al., 2008), whereas old mice continued to manifest chronic inflammation without resolution at day 68 (clinical score 1.0 ± 0.57). Mixed-effects analysis revealed a significant interaction between time and group (p < 0.001), indicating that temporal progression of clinical scores differed between young and old mice. Consistently, the cumulative inflammatory burden during the chronic phase (Day29–68), quantified as the area under the curve of longitudinal clinical scores, was significantly higher in old mice than in young mice (median value: young, 4.25 vs. old, 52.25), indicating prolonged persistence of inflammation in aged animals (Figure 1b). Based on these results, we operationally defined 2- and 5-weeks post-immunization as the acute and chronic phases for subsequent analyses, and present representative fundus and histological images from young and old mice at these time points (Figure 1c,d).

**FIGURE 1.**
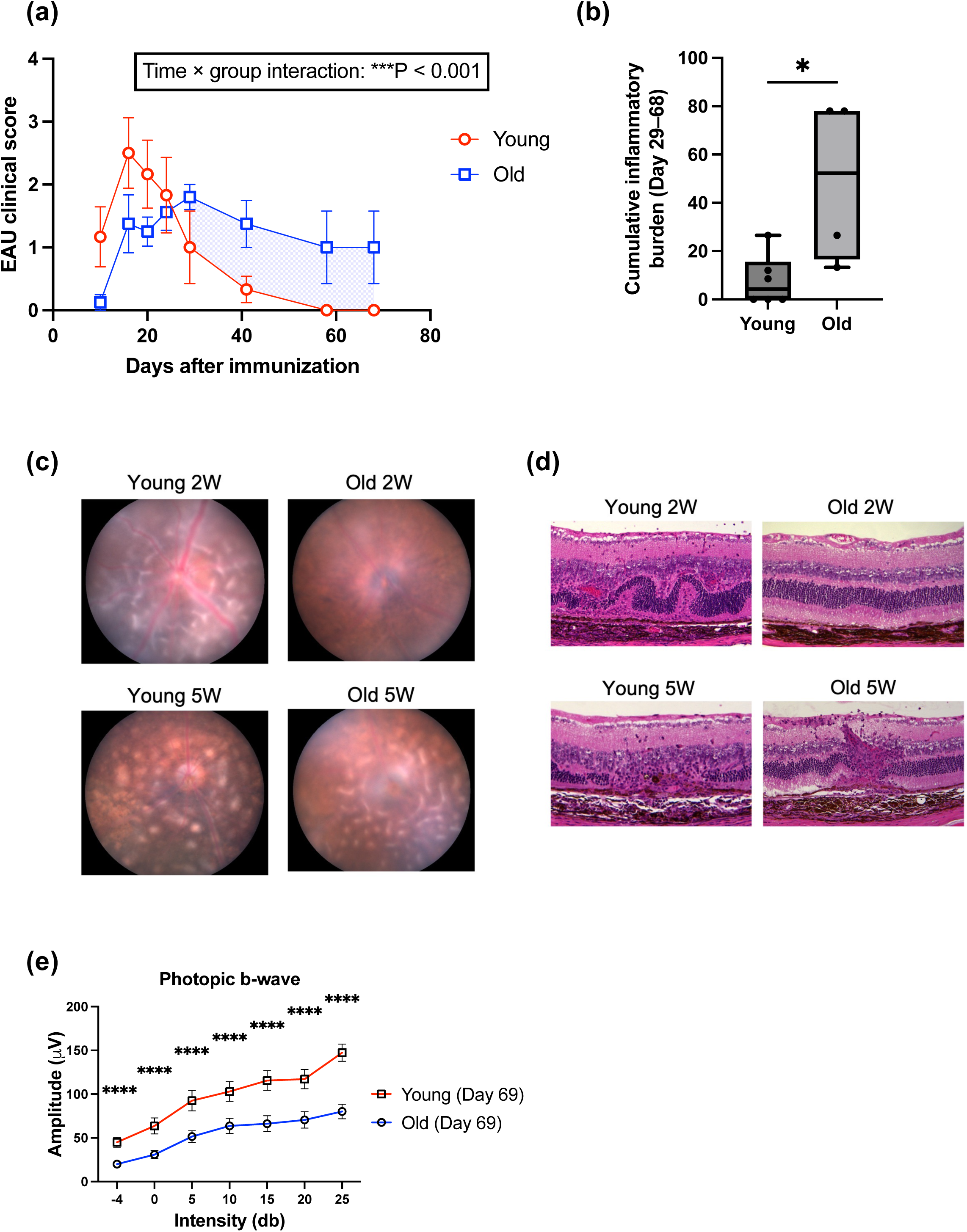
Aging alters clinical features in EAU. **(a)** Longitudinal clinical scores in young (7-week-old male, n = 6) and old mice (20-month-old male, n = 8) with EAU, analyzed by mixed-effects model, which showed a significant time × age interaction (P < 0.001). The period during which the clinical scores in old mice exceeded those in young mice (Day 29–68) is highlighted with blue shading. **(b)** The cumulative inflammatory burden was defined as the total area under the curve (AUC) of longitudinal EAU clinical scores from Day29 to Day68, reflecting sustained ocular inflammation in the chronic phase. Boxes represent the interquartile range with the median (horizontal line), and whiskers indicate the minimum and maximum values (n = 4–6). **(c and d)** Representative fundus images (c) and H&E-stained retinal sections (d) at the acute (2 weeks after immunization) and chronic (5 weeks after immunization) phases. Groups include young and old mice at each time point (Young 2W, Old 2W, Young 5W, Old 5W). **(e)** Full-field electroretinography assessing cone-driven inner retina activity (photopic b-wave), measured at day 69 (n = 8–12). *: P < 0.05. ***: P < 0.001. ****: P < 0.0001. Mixed-effects model (a). Mann–Whitney U test (b). Two-way ANOVA with Bonferroni’s multiple comparison test (e).

To assess whether sustained inflammation in old mice impacts visual function, we conducted electroretinography (ERG), which functionally assesses various retinal layers in response to light stimuli. To evaluate the long-term effects of EAU on visual function, ERG measurements were performed at baseline and at the end of the longitudinal EAU study (day 69), after the resolution of inflammation in young mice. Visual function, as assessed by scotopic a-wave, scotopic b-wave, and photopic b-wave, which reflect the function of rod photoreceptors, intermediate neurons such as bipolar cells, and cone photoreceptors, respectively, tended to be lower in old mice compared to young mice across all parameters at both baseline (pre-immunization) and the end of the study, with significant reductions observed in scotopic b-wave and photopic b-wave amplitudes at the end of the study (Figure 1e and Figure S1a,b). Notably, the photopic b-wave, which is considered to reflect cone photoreceptor function under daylight conditions, was significantly reduced at all tested light intensities in old mice both at baseline and at the end of study (Figure S1c). It has been previously reported in humans that baseline visual function assessed by ERG declines with age (Birch & Anderson, 1992; Weleber, 1981). Although the proportional decline in the ERG waveforms after uveitis was similar in young and old mice at the end of the study, the ERG at baseline in older mice translated to lower visual function at the end of the study indicating that older mice were at higher risk of vision loss given the lower baseline functional reserve capacity. To determine whether these functional changes were accompanied by retinal structural alterations, we measured retinal thickness in the major nuclear layers using optical coherence tomography. No significant changes were observed in the thickness of the inner nuclear layer, which reflects intermediate neurons, or the outer nuclear layer, which corresponds to photoreceptor cell bodies, before and after EAU in either young or old mice (Figure S1d–g), suggesting visual impairment was greater than structural loss in old mice after EAU.

These phenotypic findings showed that in old mice, the peak of EAU inflammation is delayed, and the course of inflammation is prolonged and continues without resolution. Although visual function as assessed by ERG is affected in both young and old mice, the reduced baseline visual function in old mice, as previously reported (Lin et al., 2016), potentially puts these animals at higher risk of further visual loss with chronic inflammation.

### 2.2 Macrophages and CD4^+^ T cells are the predominant immune cells in EAU retinas

To identify which immune cell populations are prominently involved in the retina during EAU, we performed comprehensive flow cytometric profiling of CD45^+^ immune cells at 2 and 5 weeks after EAU induction. Retinal cells dissociated from EAU eyes were gated on CD45 and classified into major immune populations (Figure S2a).

The number of CD45^+^ cells mirrored the clinical courses of each group. Young mice had significantly higher cell counts at 2 weeks, whereas old mice had greater numbers at 5 weeks (Figure 2a). However, there were interesting differences in the composition of this immune population over time. By analyzing the proportions of each major immune population within the total live single cells, we found that during the acute phase, young mice exhibited significantly higher proportions of most immune populations, including neutrophils, macrophages, total T cells, CD4^+^ T cells, NK cells, and NKT cells, compared to old mice (Figure 2b–f and Figure S2b–l). However, in the chronic phase, old mice developed a greater infiltration of cells suggestive of a lymphocytic response, including total T cells, CD4^+^ T cells, CD8^+^ T cells, dendritic cells, NK cells, and B cells.

**FIGURE. 2.**
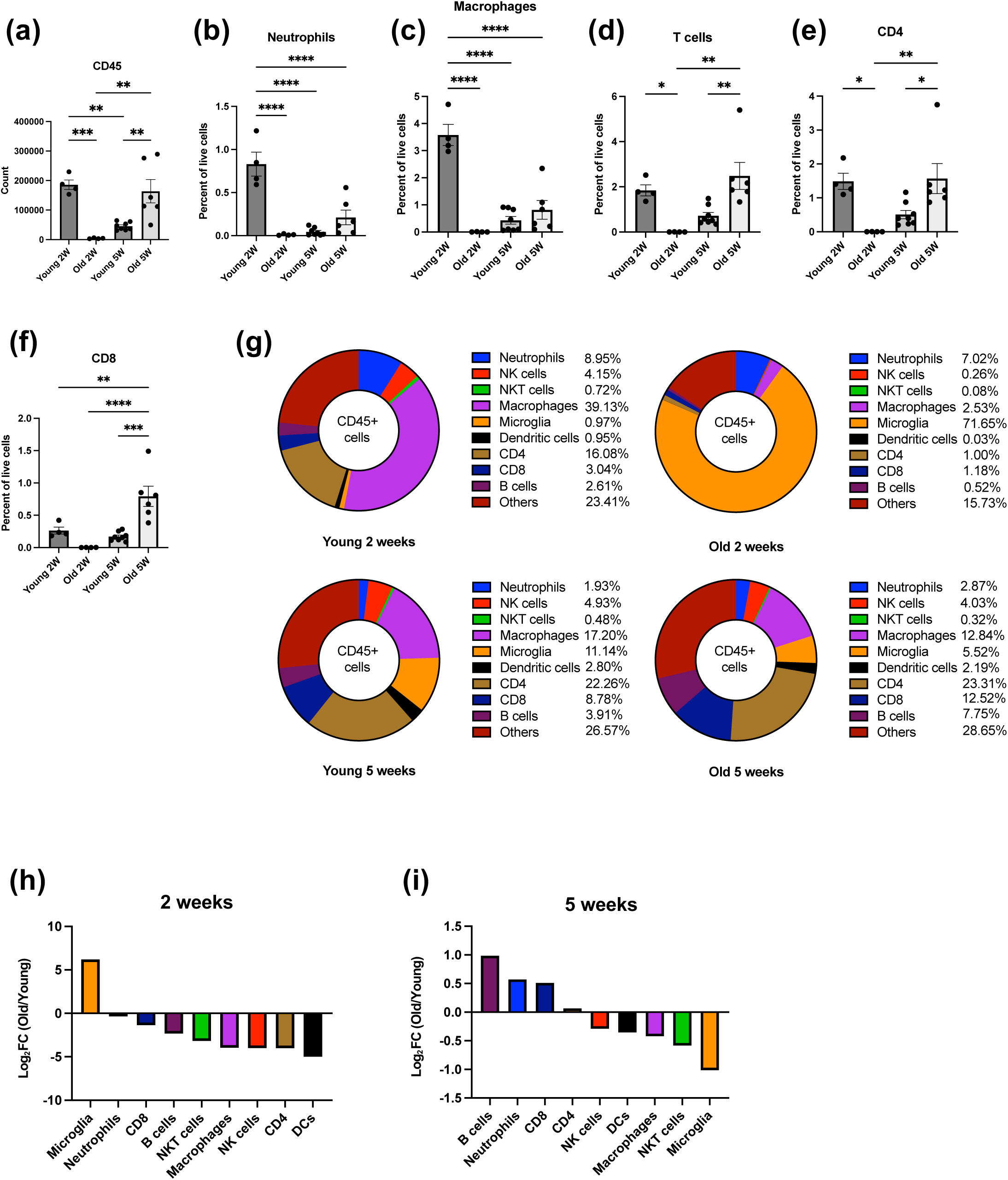
Comprehensive flow cytometric profiling of EAU retinas reveals CD4^+^ T cells and macrophages as the predominant immune cell populations. **(a–f)** Immune cell subsets in EAU retinas from young and old mice were analyzed at 2 or 5 weeks post-immunization (Young 2W, n = 4; Old 2W, n = 4; Young 5W, n = 8; Old 5W, n = 6). Total CD45^+^ cell counts (a), percentages of innate immune cell subsets among total live cells (b–f). **(g)** Immune cell compositions within CD45⁺ cells in each group. **(h and i)** Ratios of immune cell subset percentages in old versus young mice at the acute (h) and the chronic (i) phases of inflammation. Ratios were calculated by dividing the percentage in old mice by that in young mice and are presented as log_2_ fold change. *: P < 0.05; **: P < 0.01; ***: P < 0.001; ****: P < 0.0001; ns: not significant. One-way ANOVA with Tukey’s post hoc test (a–f).

Similarly, analysis of the proportion of each immune cell population within CD45⁺ cells showed that the inflammatory response consisted of CD4⁺ T cells and macrophages except for old mice during acute disease, where the immune population more closely resembled a naïve retina, with a preponderance of retinal microglia among CD45^+^ cells (Figure 2g–i). Over time, the proportion of innate immune cells decreased in the young mice between the acute and chronic phases of EAU while the lymphocytic populations increased. In contrast, old mice had an increased proportion of both infiltrating innate immune cells and even more so lymphocytes in the chronic phase, to the extent that old mice had a greater representation of lymphocytes than even young mice.

### 2.3 Age-dependent macrophage-CD4⁺ T cell interactions modulate T cell differentiation

To precisely delineate immune populations in young and old mice throughout the course of EAU, and to understand the transcriptomic basis underlying the profile differences observed between young and old mice described above, we performed scRNA-seq on retinal cells over the course of disease. Retinas were collected from young and old mice at 2 and 5 weeks after EAU induction and CD45^+^ cells were enriched by magnetic bead sorting. Cell subsets were visualized in the Uniform Manifold Approximation and Projection (UMAP) space and cell annotation was performed using canonical markers (Figure 3a and Figure S3a). Each clustered cell type was further analyzed for CD45 expression, and only CD45⁺ cells were included in the subsequent analyses (Figure 3b).

**FIGURE 3.**
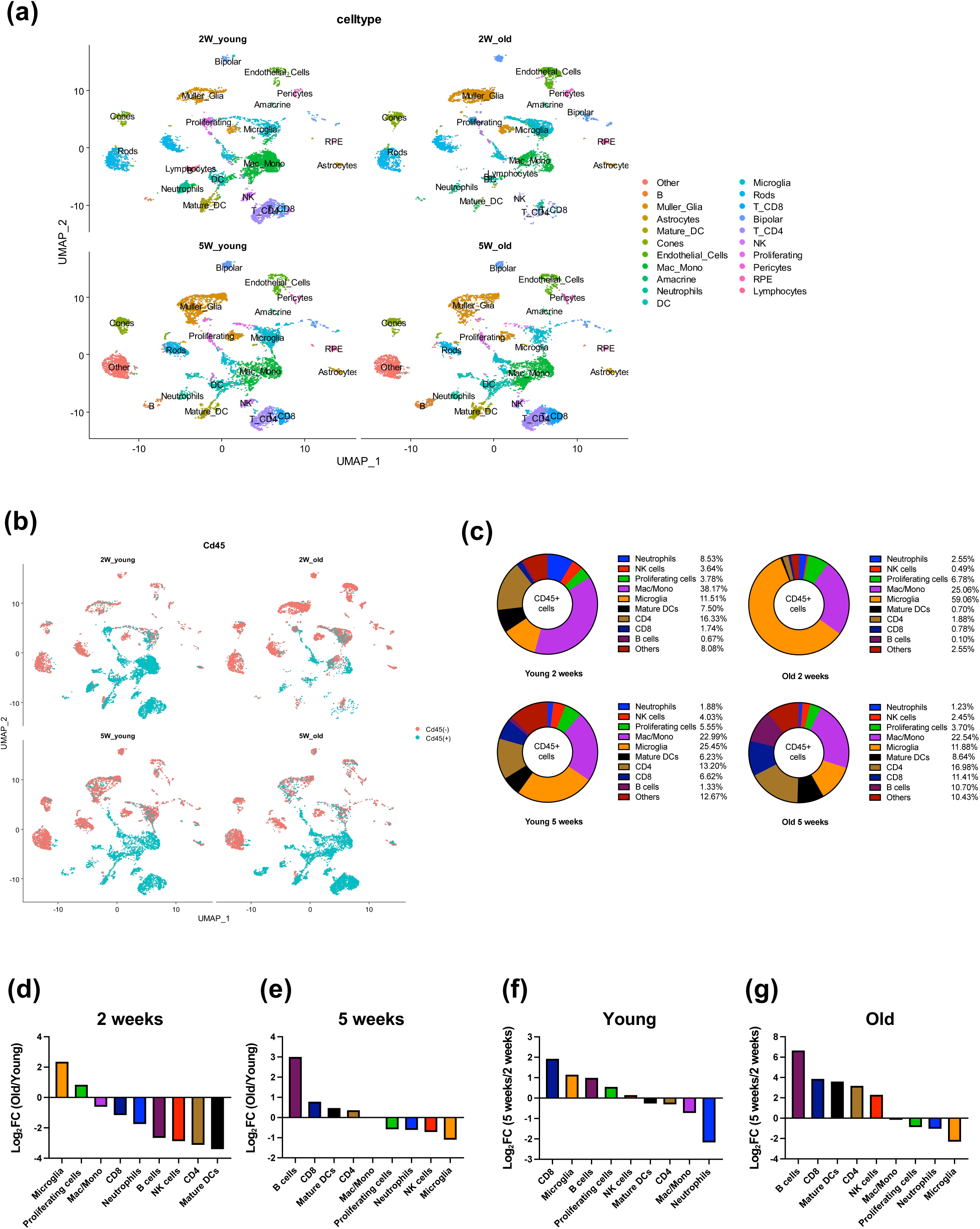
Single-cell RNA-seq of CD45^+^ retinal immune cells illustrates age- and time-dependent remodeling of the immune landscape during EAU. **(a)** UMAP plots showing immune and retinal parenchymal cell clusters in young and old EAU mice at each time point. **(b)** Re-clustering based on CD45 expression identifies immune cell populations. **(c)** Proportions of immune cell subsets within CD45^+^ cells in each group as determined by scRNA-seq. **(d–g)** Comparative analysis of immune cell subset proportions across age and disease phase. Log_2_ fold changes in immune cell subset percentages were calculated for old versus young mice at the acute phase (d), old versus young mice at the chronic phase (e), chronic versus acute phase in young mice (f), and chronic versus acute phase in old mice (g).

Quantification of cell identities in the scRNA-seq data corroborated the findings obtained by flow cytometry, with macrophages/monocytes and CD4^+^ T cells representing the predominant subsets in all groups except old mice during acute EAU (Figure 3c), where microglia remained the predominant population. Despite the persistently low proportion of CD4⁺ T cells, macrophage/monocyte clusters were markedly expanded in the old EAU group at 2 weeks, possibly due to the higher sensitivity of scRNA-seq in detecting subtle changes. Similarly, quantification of cell identities also corroborated the reversal in the proportion of microglia between young and old groups that occurred between the acute and chronic phases (Figure 3d,e) and a shift from innate to adaptive immune responses over time that was more pronounced in old mice (Figure 3f,g).

Based on the results of comprehensive flow cytometry and scRNA-seq–based cell proportion analysis, we selected peripheral immune cell subsets expected to exhibit frequent intercellular communication during both the acute and chronic phases for ligand-receptor interaction analysis. By using the tool SingleCellSignalR, known ligand-receptor pairs were analyzed to infer intercellular communication between immune cell subsets in the scRNA-seq dataset (Cabello-Aguilar et al., 2020). For this analysis, we included ligand-receptor pairs in which both the ligand and receptor genes were expressed at a value of absolute log2FC > 0.5 in either the young or old group. The resulting interactions were visualized in an alluvial diagram, where the width of each band reflects the number of ligand–receptor interaction pairs between a ligand-expressing and receptor-expressing cell type. At 2 weeks after EAU induction, macrophages represented the predominant source and recipient of cytokine signals (Figure 4a). At 5 weeks, macrophages remained the major source of cytokines; however, CD4^+^ T cells became the most predominant recipient (Figure 4b). Among all ligand–receptor interactions, macrophage-CD4^+^ T cell interaction was the most prominent at both time points.

**FIGURE 4.**
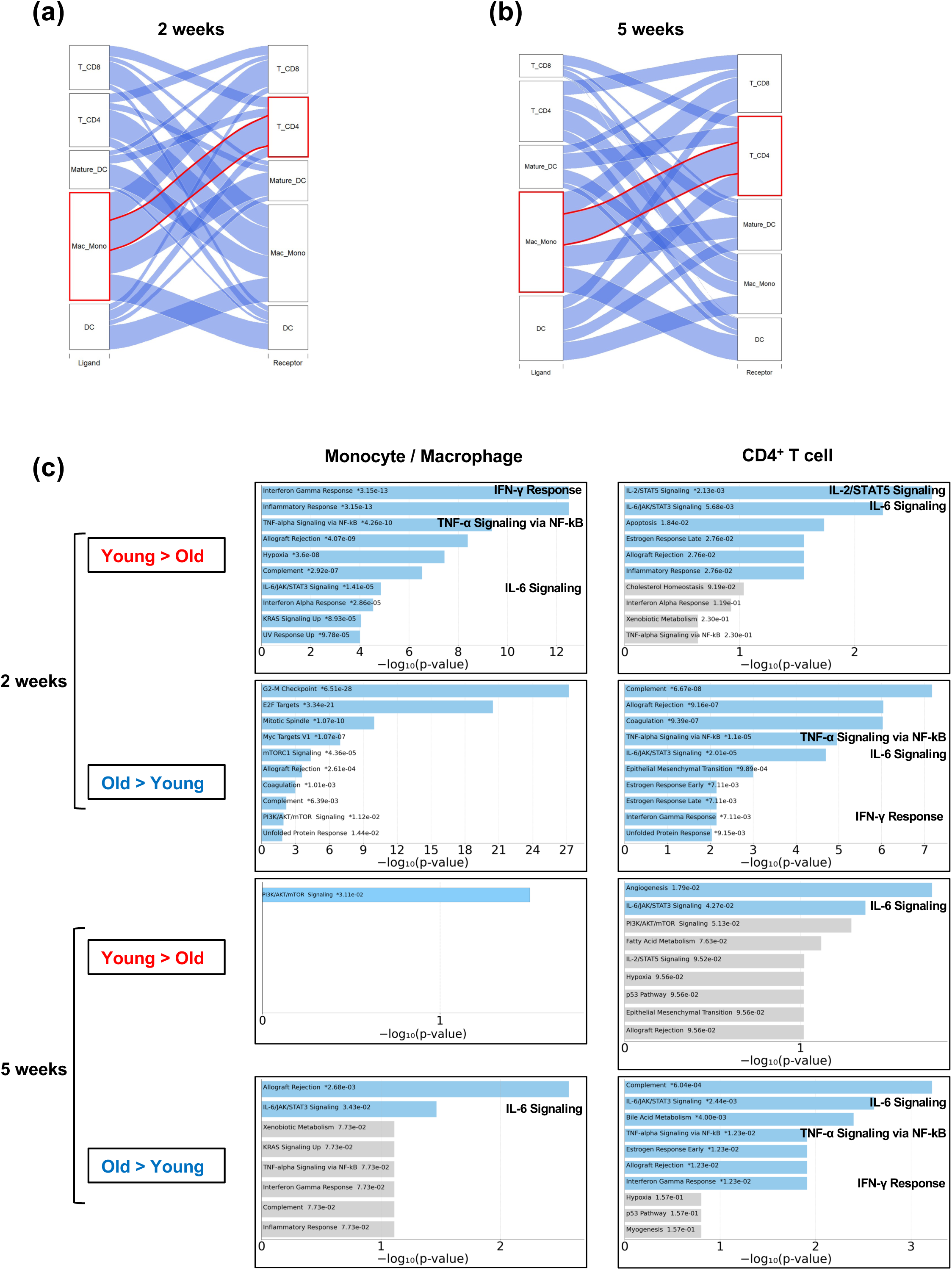
Macrophage–CD4^+^ T cell interactions shape age-dependent modulation of T cell differentiation. **(a and b)** Potential ligand-receptor pairs among dominant infiltrating immune cells were identified based on cell type specific gene expression using SingleCellSignalR. Results are visualized as alluvial plots at the acute (a) and chronic (b) phases. **(c)** Functional annotation of genes in monocyte/macrophage and CD4^+^ T cell clusters. The top 10 enriched gene sets are shown, ranked by -log_10_(P value). Bar length represents statistical significance, and gene sets with P < 0.05 are highlighted in blue. Enriched CD4^+^ T cell proliferation/differentiation-related gene sets are labeled on the right side of each graph.

According to the results of cell proportion and ligand-receptor interaction analysis, we hypothesized that macrophages and CD4^+^ T cells play a central role in the different inflammatory profiles between young and old mice with EAU. Thus, we further performed enrichment analysis by using Enrichr to identify what types of gene sets are essential for these processes (Chen et al., 2013). Statistically significant differentially expressed genes (DEGs) were extracted from the scRNA-seq dataset using a threshold of adjusted P < 0.05 and log_2_ fold change (FC) > 0.58 or < – 0.58, followed by downstream computational analyses performed with MSigDB Hallmarks 2020 gene sets (Liberzon et al., 2015). In the macrophage/monocyte population, multiple gene sets related to acute inflammation were significantly enriched in young mice at the 2-week time point including interferon-γ (IFN-γ) response, tumor necrosis factor-α (TNF-α) signaling, and IL-6 signaling (Figure 4c). In contrast, there were no inflammatory cytokine signaling pathways that were upregulated in young mice at 5 weeks, and the IL-6 signaling pathway had emerged as a key difference between young and old mice with chronic uveitis. Similarly, in the CD4^+^ T cell population, the IL-6 signaling pathway was differentially regulated between young and old mice at all time points, suggesting that IL-6 signaling could be a key player in the aged immune response. Taken together, these analyses of scRNA-seq demonstrate that macrophage-CD4⁺ T cell interactions are key factors in age-dependent changes in EAU retinas and that altered CD4^+^ T cell differentiation-related pathways, especially IL-6 signaling, are dominant immunological features in old mice during disease progression.

### 2.4 In co-culture, aged macrophages promote CD4^+^ T cell proliferation while simultaneously reducing Treg frequency

We previously demonstrated that macrophage senescence induced the upregulation of proinflammatory cytokines, a phenomenon known as the senescence-associated secretory phenotype (SASP) (Terao et al., 2024). To assess the impact of macrophage aging on macrophage–CD4^+^ T cell interactions in old mice, we performed a co-culture assay using CD4^+^ T cells isolated from young OT-II mice and bone marrow-derived macrophages (BMDMs) from either young or old mice (Figure 5a). Equivalent numbers of BMDMs were seeded and stimulated with lipopolysaccharide, followed by the addition of CFSE-labeled CD4^+^ T cells at a 1:1 cell ratio. T cell proliferation was determined by CFSE dilution, visualized as multiple peaks by flow cytometry. CD4^+^ T cells showed significantly greater proliferation when co-cultured with aged BMDMs compared to young BMDMs (Figure 5b,c). We further analyzed the concentrations of multiple cytokines secreted into the supernatants of these co-cultures using a Luminex assay and ELISA. Among the cytokines critically involved in CD4^+^ T cell differentiation and activation, IL-2, IL-6, and IL-17 were elevated in the aged macrophage co-culture, with IL-6 showing the most striking increase (Figure 5d,e and Figure S4a). In contrast, TGF-β, which is essential for the differentiation of Treg, showed no significant difference between the two groups (Figure 5f). To identify the cellular sources of these cytokines, we assessed mRNA expression in BMDMs after LPS stimulation and in CD4^+^ T cells isolated by FACS after 72 hours of co-culture. Aged BMDM showed higher expression of IL-6 mRNA at 3 and 6 hours after stimulation (Figure 5g), but we found no significant difference in IL-6 expression in co-cultured CD4^+^ T cell population (Figure S4b). Since IL-6 plays a crucial role in directing the fate of naïve CD4^+^ T cells and whether they acquire a Treg phenotype (Kimura & Kishimoto, 2010), we hypothesized that aged macrophages are deficient in differentiating naïve CD4^+^ T cells into Tregs. In order to test this hypothesis, we re-clustered CD4^+^ T cells from our scRNA-seq dataset and found a trend toward a higher proportion of Tregs in old mice compared to young mice during both the acute and chronic phases, although statistical power was limited due to the sample size and the relatively small number of recovered Treg cells (Figure S3b,c). Analysis of the Treg/BMDM co-culture revealed a significant decrease in the proportion of Foxp3^+^ Tregs among CD4^+^ T cells co-cultured with aged BMDMs compared to those co-cultured with young BMDMs (Figure 5h,i).

**FIGURE 5.**
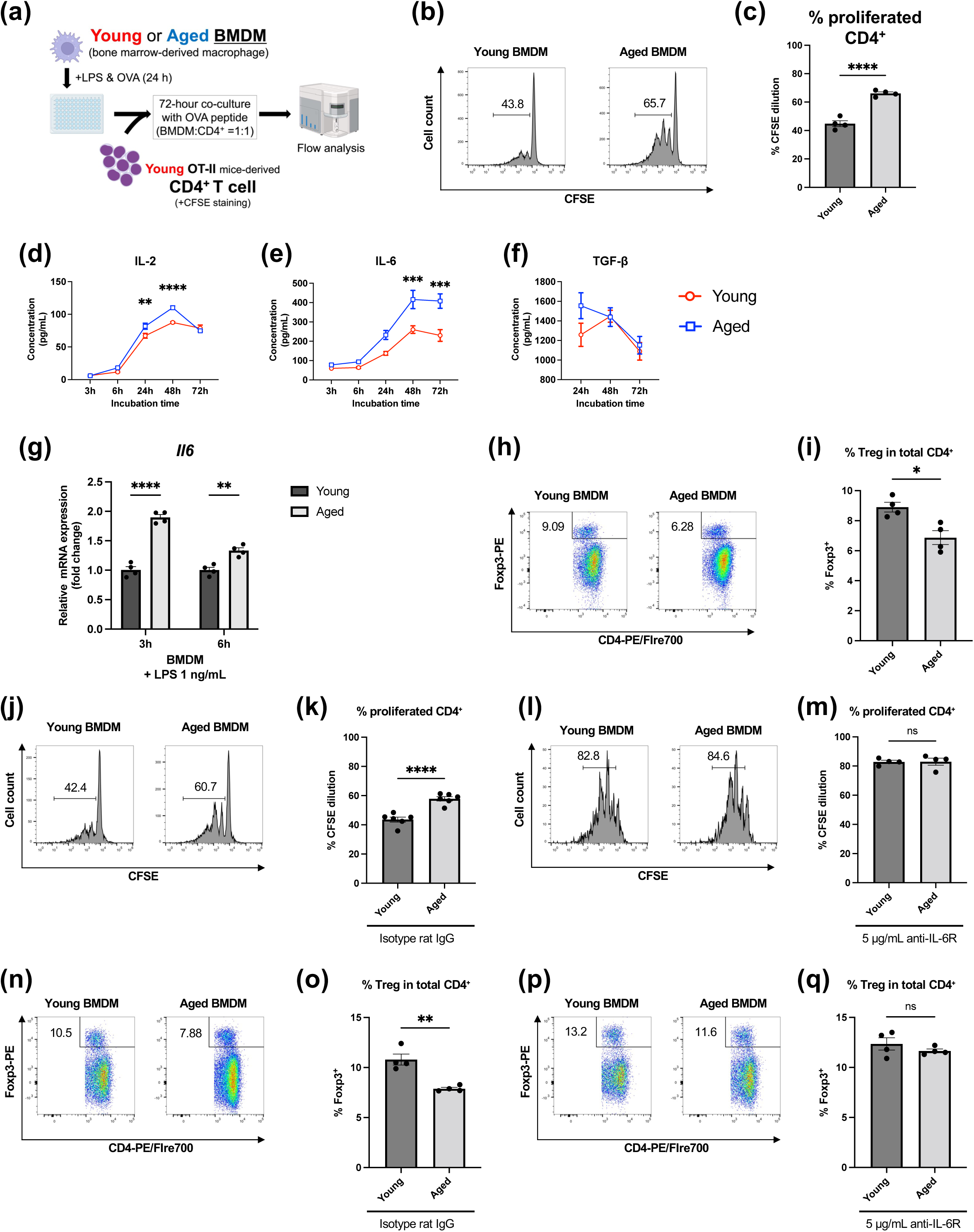
Co-culture assays of CD4^+^ T cells with BMDMs demonstrated that aged BMDMs promote IL-6–dependent T cell proliferation while impairing Treg induction. **(a)** Schematic of the co-culture assay using bone marrow-derived macrophages (BMDMs) and CD4^+^ T cells. **(b and c)** Proliferation of CFSE-labeled CD4^+^ T cells derived from young OT-II mice in a co-culture assay with BMDMs at a 1:1 ratio (n = 4). **(d–f)** Cytokine concentrations of IL-2 (d), IL-6 (e), and TGF-β (f) in co-culture supernatants (n = 3–4). **(g)** Gene expression of IL- 6 in BMDMs after 3 and 6 hours of LPS stimulation (n = 4). **(h and i)** Frequencies of Foxp3^+^ cells among CD4^+^ T cells in BMDM co-cultures (n = 4). **(j–m)** Proliferation of CFSE-labeled CD4^+^ T cells treated with isotype control antibody (j and k) or anti-IL-6 receptor antibody (l and m) (n = 4–6). **(n–q)** Frequencies of Foxp3^+^ cells among CD4^+^ T cells in BMDM co-cultures treated with isotype control antibody (n and o) or anti-IL-6 receptor antibody (p and q). *: P < 0.05; **: P < 0.01; ***: P < 0.001; ****: P < 0.0001; ns: not significant. Unpaired Student’s t test (c, g, i, k, m, o, and q). Two-way ANOVA with Bonferroni’s post hoc test (d–f).

To determine whether the enhanced T cell proliferation and reduced Treg proportion observed in the aged BMDM co-culture might be mediated by IL-6, we inhibited IL-6 signaling using an IL-6 receptor–neutralizing antibody. While incubation with isotype control antibody had no effect on the increased CD4^+^ T cell proliferation when cultured with aged BMDMs, this difference was abolished by IL-6 receptor blockade (Figure 5j–m). Similarly, IL-6 receptor blockade also abolished the differences in Foxp3^+^ Treg induction between young and aged BMDM co-culture (Figure 5n–q). These findings indicate that IL-6 signaling is essential for the aged BMDM-induced enhancement of CD4^+^ T cell proliferation and a simultaneous reduction in the proportion of Treg.

### 2.5 Old mice demonstrate a reduction in induced Treg frequency during EAU

Foxp3^+^ Tregs are generally classified into thymus-derived Tregs (tTregs) that differentiate during T cell development and peripherally induced Tregs (pTregs) that differentiate in the periphery from naïve CD4^+^ T cells (Yadav et al., 2013). Markers such as Helios (Ikzf2) have been proposed to help distinguish these subsets (Singh et al., 2015). A previous study suggested that inflammation in EAU is suppressed not by tTregs but by induced Tregs (Lee & Taylor, 2015), and depletion of Foxp3^+^ Tregs impaired the ability to resolve EAU-mediated inflammation and induced relapses during the remission phase (7–12 weeks post-induction), indicating their long-term role in suppressing recurrent intraocular inflammation (Silver et al., 2015). Therefore, we investigated whether the age-related immune changes in EAU, in particular the persistence of inflammation, could be due to an impaired induction of Treg in the periphery in old mice. We examined the proportion of pTregs in the cervical draining lymph nodes (CDLNs), which are essential organs for antigen sensing and regulating intraocular immune responses (Yin et al., 2024). To account for the time lag between inflammatory changes and their impact on Treg differentiation in the CDLNs, we harvested CDLNs from young and old mice at 3 and 6 weeks after EAU induction, corresponding to the post-acute phase and post-chronic phase, respectively. In the post-acute time point, there was no difference between tTreg and pTreg (Figure 6a–c). In contrast, the post-chronic time point demonstrated a shift in the Treg populations in the CDLN, with a decrease in Helios^-^pTregs in old mice and a corresponding increase in tTreg (Figure 6d–f). These findings mirror the *in vitro* Treg dynamics observed in the co-culture assay and support the notion that pTreg induction is reduced in old mice, especially during the chronic phase.

**FIGURE 6.**
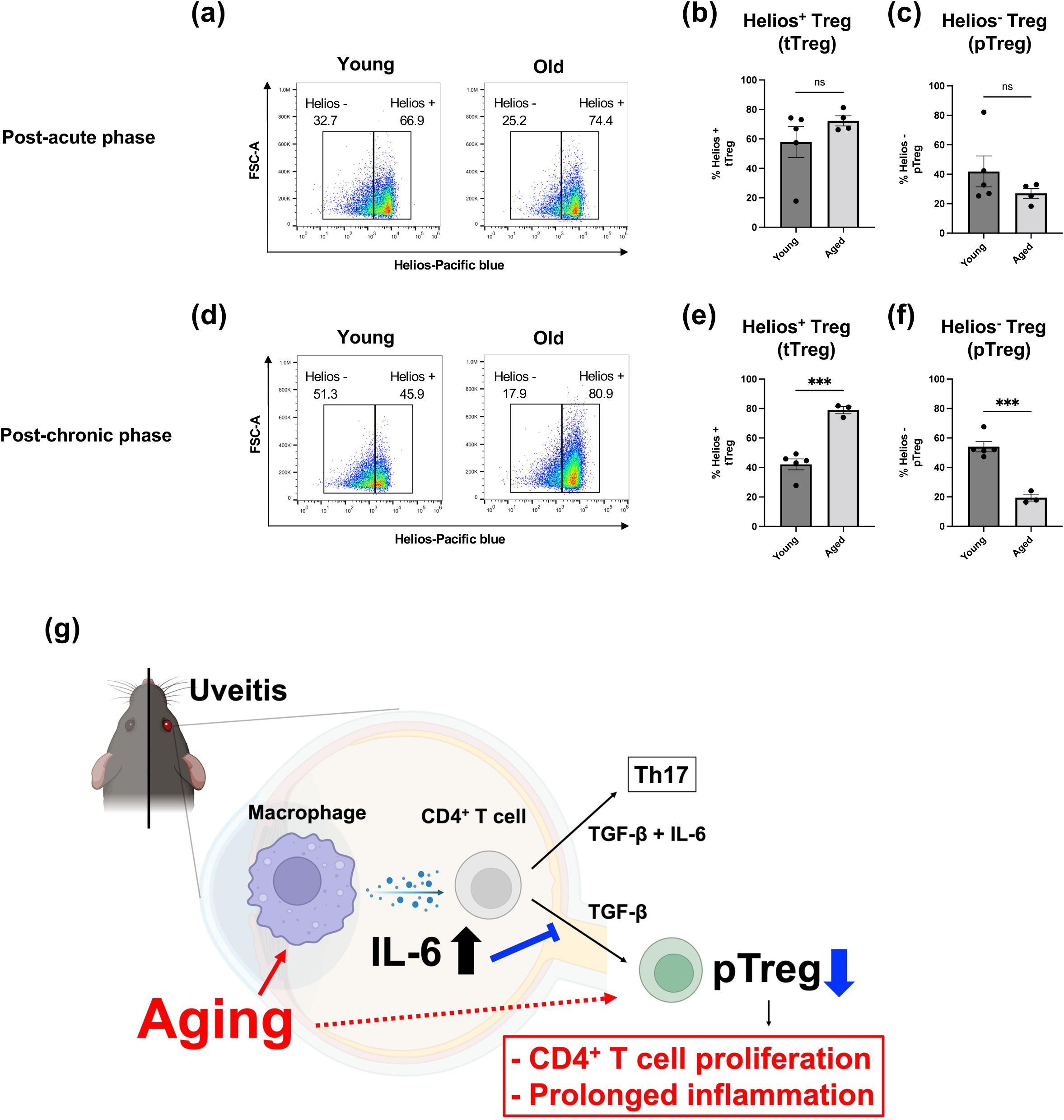
Treg induction is suppressed at the chronic phase of EAU in old mice. **(a–f)** Frequencies of Helios^+^ thymus-derived Treg (tTreg) and Helios^-^ peripherally induced Treg (pTreg) in the draining cervical lymph nodes. Comparisons of young and old mice at 3 weeks (a–c) and 6 weeks (d–f) after EAU induction (n = 3–5). **(g)** Schematic summary of results. Aged macrophages shift the CD4^+^ T cell differentiation balance toward Th17 over Treg via IL-6, leading to increased T cell proliferation and prolonged inflammation by reduced pTreg induction. ***: P < 0.001; ns: not significant. Unpaired Student’s t test (b, c, e, and f).

Taken together, these results demonstrate that aging is associated with a delayed initial immune response to antigen in the EAU model, but once initiated, old mice are unable to efficiently dampen the effector immune response. This leads to a persistent, chronic inflammatory phenotype in old mice that is associated with impaired visual function. Mechanistically, our study supports that increased macrophage IL-6 secretion contributes to greater CD4^+^ T cell proliferation and a reduction in Helios^-^ pTregs that are critical in dampening EAU.

## 3 Discussion

Inflammaging or immunosenescence is implicated in age-associated dyshomeostasis and in the pathogenesis of diverse age-related diseases including cancers, cardiovascular diseases and age-related eye diseases (Liu et al., 2023; Terao et al., 2024; Wang et al., 2019). Non-infectious uveitis, characterized by immune-mediated intraocular inflammation, is a leading cause of blindness both globally (Maghsoudlou et al., 2025) and in the US (Rothova et al., 1996). Advanced age is a well-known factor that affects the presentation and clinical course of inflammation in uveitis (Samalia et al., 2025), with a much higher proportion of vision loss in older individuals compared to younger counterparts (Maini et al., 2004). Despite this, the age-associated alterations that lead to this increased vision loss are unknown. Immunosenescence, the alteration in the immune system with aging, has significant effects on both the adaptive and innate immune system. While T cells are known to be the primary drivers of autoimmune uveitis, disease pathology is heavily influenced by the immunologic microenvironment (Kerr et al., 2008). Macrophages, in particular, play a vital role in the immune response as both downstream effectors of T cell responses and as signaling partners that can both activate and modulate T cell responses (Merida et al., 2015). Here, using multimodal analyses *in vitro* and *in vivo*, we demonstrated a central role for abnormal macrophage-CD4^+^ T cell interactions driving the phenotype of aging-associated uveitis. Specifically, macrophages derived from old mice significantly enhanced CD4^+^ T cell proliferation while concurrently reducing the generation of Tregs through aberrant IL-6 signaling. These results highlight a key role for inflammaging in ocular inflammation, where increased levels of IL-6, a typical SASP factor, were associated with an age-associated decrease in Treg generation. This age-associated immunosenescence, combined with increased CD4^+^ T cell proliferation, manifested as sustained, chronic inflammation in old mice (Figure 6g). Although Il-6 is known to play a key role in regulating uveitic inflammation, its contribution to the pathogenesis of uveitis associated with aging has not been explored previously.

As the primary immune cell in the homeostatic retina, and one of the primary effector immune cells in EAU (Merida et al., 2015; Sonoda et al., 2003), we explored the role of aged macrophages in the immunosenescence process. Aged macrophages are known to mediate altered CD4^+^ T cell differentiation and activation through both cytokine production and direct cell–cell contact (Lorenzo et al., 2022). Unlike acquired immune cells that demonstrate immunological memory and can persist in the long-term in the periphery, innate immune cells derived from hematopoietic stem cells acquire aging-associated profiles due to an altered bone marrow environment, which becomes myeloid-biased with stem cell aging (Geiger et al., 2013; Ross et al., 2024; Yang & de Haan, 2021). With aging, macrophages, cells that are key components of the innate immune system, acquire SASP characteristics through the process of immunosenescence, and have been previously shown to be a pivotal source of pro-inflammatory cytokines in diverse tissues including the eye (Droho et al., 2021; Hou et al., 2024; Terao et al., 2024). Consistent with age-related increases in IL-6 expression reported in both systemic and ocular tissues (Maggio et al., 2006; Terao et al., 2022), aged macrophages displayed increased IL-6 signaling, similar to the *in vivo* signaling pathways that were found to become activated during EAU in old mice. This process is likely driven by epigenetic changes in macrophage progenitors, but the exact changes leading to this proinflammatory skewing need further elucidation.

To explore the relevance of the myeloid aging phenotype to autoimmune disease, we utilized young or aged macrophages co-cultured with young responder CD4^+^ T cells to isolate the effects of the myeloid compartment during EAU. We focused on the ability of macrophages to induce regulatory CD4^+^ T cells, a vital player in preventing auto-immune effector responses. Tregs can be produced in the thymus or develop in the periphery (Chen et al., 2003; Fontenot et al., 2003), and while they demonstrate several overlapping functions, they retain distinct functionality based on their ontogeny (Shevach & Thornton, 2014; Thornton et al., 2019), with pTregs responding to microenvironmental cues, while tTregs play a larger role in controlling natural self-reactivity. These vital homeostatic cells are prone to multiple aging-related changes. Although aging increases Treg frequency and preserves their suppressive potency *in vitro* (Lages et al., 2008), which may contribute to the delayed and blunted early inflammatory peak in old mice, the resolution of chronic inflammation in EAU depends chiefly on induced Tregs (Lee et al., 2015).

This is consistent with a recent report which showed that old mice exhibited reduced Th17 responses and attenuated inflammation during the acute phase of EAU; however, that study did not address the mechanisms during the chronic phase (Li et al., 2022). Aged macrophages are markedly less efficient at converting naïve CD4^+^ T cells into this subset, creating a deficit in context-dependent pTreg induction that ultimately leads to chronic inflammation. For instance, in the experimental autoimmune encephalomyelitis (EAE) model, which shares core pathogenic mechanisms with EAU, Tregs played a crucial role in the resolution phase of inflammation, with aged Tregs having an impaired regulatory capacity (de la Fuente et al., 2024). These Treg alterations with aging contribute to the complex immune dysregulation associated with immunosenescence (Guo et al., 2020; Rocamora-Reverte et al., 2020; Yang et al., 2024) and may explain why aged animals display an impaired ability to control novel inflammatory stimuli over time.

Antigen specificity is critical for Tregs to control inflammation in EAU (Chen et al., 2021). Indeed, in EAE, which is a neuroinflammatory model of the central nervous system (CNS), old mice show impaired inflammatory control, especially in the chronic phase, that is associated with reduced representation of antigen-specific Tregs in the CNS (Wang et al., 2022). Tregs from IL-6 knockout mice show stronger IRBP-specific responses than those from wild-type mice (Haruta et al., 2011). Since Tregs need ongoing TCR stimulation to remain functional (Vahl et al., 2014), and retinal antigens are rarely presented unless the blood–retinal barrier (BRB) is breached, antigen-specific tTreg activity likely declines with age. Meanwhile, pTregs generated in response to BRB disruption may retain stronger retinal antigen specificity and suppressive capacity (Sakaguchi et al., 2023; Selck & Dominguez-Villar, 2021). Aging likely impairs both pTreg induction and tTreg function, weakening inflammation control in immune-privileged sites.

Aging is correlated with an increased incidence of vision loss in uveitis patients (Maini et al., 2004). Uveitic macular edema (UME), associated with increased vascular permeability in the central part of the retina responsible for the most precise vision, is a leading cause of uveitis-related vision loss (Lardenoye et al., 2006; Yu et al., 2024), and both aging and chronic inflammation were recognized as risk factors for its development (Accorinti et al., 2019). Meanwhile, lower serum IL-6 levels and a higher proportion of Tregs in peripheral blood were associated with better prognosis in UME (Matas et al., 2020), suggesting that IL-6, Treg activity, and UME may act in concert and contribute to vision loss in the elderly. IL-6 targeted therapies are currently used primarily for the treatment of uveitis associated with rheumatic diseases, such as rheumatoid arthritis and juvenile idiopathic arthritis (Karkhur et al., 2019). However, intravitreal administration of IL-6 antagonists has been shown to improve clinical scores in EAU, and systemic IL-6 inhibition has demonstrated efficacy in treating UME in patients with uveitis (Tode et al., 2017; Yang et al., 2023). Based on these findings, we suggest that modulation of age-associated inflammatory pathways, including IL-6 signaling, may represent a therapeutic avenue in uveitis. Collectively, our findings demonstrate that macrophage-derived IL-6, a canonical SASP cytokine, limits the differentiation of naïve CD4⁺ T cells into pTregs and thereby contributes to T cell proliferation and persistent ocular inflammation. This finding provides mechanistic insight into the immunopathogenesis of vision-threatening uveitis in the elderly and highlights the potential of targeting IL-6 signaling as a therapeutic strategy to ameliorate uveitis-associated inflammation, especially in older individuals that are much more likely to bear the brunt of vision loss from this devastating condition.

## 4 Materials and Methods

### 4.1 Mice

Young (6–12 weeks) and aged (20–21 months) male C57BL/6J mice and OT-II mice (stock no. 004194) were obtained from The Jackson Laboratory and maintained under specific pathogen–free conditions. All procedures were approved by the Washington Univsersity in St. Louis IACUC and performed in accordance with the Washington University School of Medicine Animal Care and Use guidelines, as well as the ARVO Statement for the Use of Animals in Ophthalmic and Vision Research.

### 4.2 EAU immunization and evaluation

The human interphotoreceptor retinoid-binding protein peptide sequence 651–670 (hIRBP_651–670_, LAQGAYRTAVDLESLASQLT) was synthesized by GenScript. Complete Freund’s adjuvant (CFA) and desiccated *Mycobacterium tuberculosis* strain H37Ra were purchased from Becton Dickinson and Company. Purified *Bordetella pertussis* toxin (PTX) was purchased from EMD Millipore Corporation. EAU induction was performed as previously described (Hase et al., 2021). The clinical severity of retinal inflammation and retinal thickness were assessed by fundoscopic examination using a Micron III fundus camera (Phenix Research Laboratories) and optical coherence tomography (Phenix Research Laboratories, OCT2), respectively. Mice were anesthetized and pupils were dilated using an eye drop solution of 1% tropicamide (Somerset Therapeutics). The clinical scoring was based on the number or extent of vasculitis foci and exudate in the retina, and graded on a five-point scale, as previously reported (Harimoto et al., 2014).

To measure thickness of the INL and ONL, OCT images were analyzed by ImageJ software (National Institutes of Health), and layer thickness was quantified every 250 µm from optic nerve head.

### 4.3 Electroretinogram (ERG)

Dark adopted ERG was performed as previously described (Terao et al., 2024). A UTAS BigShot System (LKC Technologies Inc.) was used to perform the measurement. The stimulus consisted of a full-field white light flash (10 µs). Raw data were processed using MATLAB software (MathWorks). The amplitude of the a-wave was measured from the average pretrial baseline to the most negative point of the average trace, and the b-wave amplitude was measured from that point to the highest positive point.

### 4.4 Flow cytometry

Bilateral retinas from each mouse were pooled and considered as a single sample for assessment in comprehensive flow cytometry. Each sample was dissociated using Collagenase D and Collagenase type VIII, followed by resuspension in FACS buffer and Fc receptor blocking with DAPI staining. For characterization of immune subsets, antibodies listed in Table S1 were used at a 1:50 dilution in FACS buffer. Flow cytometry data were acquired using Cytek Aurora (Cytek Biosciences). For the Treg analysis in cervical draining lymph nodes (CDLN), CDLNs collected from young and old mice at 3 or 6 weeks after EAU induction were mashed on a 40 µm cell strainer with FACS buffer. CDLN cell suspensions were stained with LIVE/DEAD Fixable Near-IR Dead cell staining kit (Thermo Fisher Scientific) and antibodies for Treg analysis as detailed in Table S1. All antibodies were used at a 1:100 dilution in FACS buffer. Flow cytometry data were acquired using Attune NxT Flow Cytometer (Thermo Fisher Scientific). All the data were analyzed using FlowJo v10 software (BD Biosciences).

### 4.5 Fluorescence-activated cell sorting (FACS)

FACS was carried out with a MoFlo Cell Sorter (Beckman Coulter) at the Siteman Cancer Center Flow Cytometry core. CD4^+^ cells were identified as PE/Fire700-positive and DAPI-negative, and sorted directly into RLT lysis buffer (Qiagen). Samples were snap-frozen and stored at -80 ℃ until RNA isolation.

### 4.6 Single cell RNA sequencing (scRNA-seq)

We conducted single cell RNA sequencing of mouse retina in EAU using the 10x Genomics platform. We prepared single cell suspensions of mouse retina using a papain-based digestion (Pfeifer et al., 2023). For each experimental group, we pooled neural retinas from n = 4–6 mice and enriched the single cell suspension for CD45^+^ cells using the EasySep Mouse CD45 Positive Selection Kit (STEMCELL Technologies). Finally, we washed and resuspended the cells in DMEM media containing 10% fetal bovine serum and used them as input for single cell RNA sequencing using the 10x Genomics Chromium Single Cell 3’ v3.1 Reagent Kit as previously described (Lin et al., 2023). We sequenced the samples on the Illumina NovaSeq 6000 platform at the McDonnell Genome Institute at Washington University School of Medicine in St. Louis.

### 4.7 Single cell RNA sequencing computational analysis

First, we processed the raw FASTQ sequencing files using CellRanger 6.1.1 with alignment to the 10x Genomics mouse reference genome (refdata-gex-mm10-2020-A). We imported the filtered count matrices into Seurat v5 (Hao et al., 2024), and assigned each cell a unique identifier to prevent overlap of barcodes between different samples. We normalized, log-transformed, and scaled the count matrices to remove unwanted sources of variation such as discrepancies in sequencing depth. We identified the top 2000 highly variable genes for principal component analysis. We integrated across samples to account for any sample- or experiment-specific batch effects using the Harmony package (Korsunsky et al., 2019). We utilized these harmony embeddings to run UMAP dimensional reduction to 2 dimensions. We clustered cells according to the Louvain algorithm and identified marker genes for each cluster using Wilcoxon rank sum tests comparing each cluster to all other clusters. Marker genes for each cell type are provided in Supplementary Fig. 3 A.

For re-clustering of Tregs, we first subsetted CD4^+^ T cells and processed them using the same pipeline, including batch correction with Harmony. After UMAP visualization and clustering in the Harmony space, we identified a cluster enriched for canonical Treg markers (*Foxp3*, *Il2ra*, *Ctla4*, and *Cd4*), which was designated as the Treg population.

To identify ligand-receptor signaling dysregulated in aging, we used the package SingleCellSignalR (Cabello-Aguilar et al., 2020). We first generated the full signaling network with the function cell_signaling() and the following parameters: int.type = ‘paracrine’, s.score = 0.5, and tol = 0.05. To narrow the full signaling network to those that were dysregulated in aging, we filtered out interactions for which either the ligand and/or receptor was dysregulated in aging (logfc.threshold = 0.5, p_val_adj < 0.05). To focus on consistently abundant infiltrating immune populations across groups, we restricted the analysis to five immune cell types (DC, Mac_Mono, T_CD4, T_CD8, and Mature_DC), excluding tissue-resident microglia, B cells, neutrophils, NK cells, proliferating cells, and retinal parenchymal cells.

DEGs were defined for each cell type at each time point based on an absolute log_2_ fold change > 0.58 and an adjusted P value < 0.05, as determined from scRNA-seq data set. DEGs were then subjected to pathway enrichment analysis using Enrichr (Chen et al., 2013), with the MSigDB Hallmark gene sets (2020 version) used as the reference database.

### 4.8 Bone marrow-derived macrophages and CD4^+^ T cells co-culture assay

Bone marrow-derived macrophages (BMDMs) were isolated as previously described (Terao et al., 2024). Briefly, 2- or 20-month-old male mice were euthanized, and bone marrow cells were isolated from limb bones by flushing with DMEM media. Isolated cells were cultured in a humidified incubator at 37 ℃ with 5% CO_2_ with DMEM media containing 10% conditioned media collected from CMG 14-12 cells containing macrophage colony stimulating factor for 6 days. Following incubation, macrophage-differentiated cells were harvested using Accutase and subsequently stored in liquid nitrogen.

Thawed BMDMs were seeded at 1 × 10^5^ cells per well in 96-well round-bottom plates and cultured in DMEM supplemented with 10% FBS, 1 ng/mL LPS, and 2 µg/mL ovalbumin peptide (OVA_323–339_; AnaSpec). After 24 hours, the cells were washed with PBS and the medium was replaced with T cell complete medium. Spleens and lymph nodes were harvested from 5- to 8-week-old OT-II mice, mechanically dissociated through a 40 µm cell strainer and resuspended in FACS buffer. CD4⁺ T cells were then isolated using the EasySep Mouse CD4^+^ T Cell Isolation Kit (STEMCELL Technologies) according to the manufacturer’s protocol. For the T cell proliferation assay, isolated CD4^+^ cells were labeled with 2.5 µM CFSE (Tonbo Biosciences) according to the manufacturer’s protocol. CD4^+^ T cells were co-cultured with BMDMs at a 1:1 cell ratio in T cell complete medium containing 2 µg/mL OVA for 72 hours. Cells were then stained with CD4-PE/Fire700, FOXP3-PE, TCRβ-APC, and DAPI (antibody details are shown in Table S1), and analyzed using Attune NxT Flow Cytometer.

### 4.9 Cytokine quantification

Cytokine levels (except for TGF-β) in the cell culture supernatants from the co-culture assay were measured using the Mouse Luminex Discovery Assay (R&D Systems). The samples were stored at -80℃ until being thawed on ice, then 50 microliters of 2-fold diluted sample or prepared kit standard was added to each well (in duplicate) of a 96 well plate containing premixed beads. The matrix for the standard and background wells was base culture media. Calibrator diluent (RD6-52) was used to dilute the samples. The bead-based multiplex immunoassay was performed according to the manufacturer’s instructions (R&D Systems, LXSAMSM-17). The bead MFI was measured using a FLEXMAP3D Luminex (Luminex Corp) machine. A dilution factor of 2 was applied to each sample well value in the Luminex xPONENT42 software. Analysis software, MilliporeSigma Belysa v.1.2.1 (Merck EMD Millipore), was used to calculate the pg/ml for each analyte using a 5-parameter logistical curve-fit algorithm.

For quantification of TGF-β, supernatant samples were activated using the Sample Activation Kit 1 (R&D systems), followed by measurement with the Mouse TGF-beta 1 DuoSet ELISA (R&D systems) per the manufacturer’s instructions. Optical density was determined using a Spark multimode microplate reader (Tecan).

All samples were analyzed in duplicate for both the multiplex cytokine assay and the ELISA, and mean values were used for subsequent analysis.

### 4.10 RNA isolation and reverse transcription qPCR

Total RNA was isolated from FACS-sorted cells using the RNeasy Micro Kit (Qiagen) and from all other samples using TRIzol, followed by RT-qPCR as previously described (Terao et al., 2024). qPCR was performed using TaqMan gene expression assays for *Il6* (Mm00446190_m1) and *Actb* (Mm02619580_g1) with TaqMan Fast Advanced Master Mix on a StepOnePlus Real-Time PCR System (Thermo Fisher Scientific). *Actb* was utilized as the internal control and relative expression levels were calculated using the ΔΔCt method.

### 4.11 Statistics

All results are expressed as mean ± SEM. Student’s *t*-test was used for statistical comparison between groups, and one-way analysis of variance (ANOVA) followed by the Tukey–Kramer method as a post hoc test was used for multiple comparison procedures. For time-course cytokine measurements between two groups, two-way ANOVA with Bonferroni’s post hoc test was performed. Longitudinal clinical scores were analyzed using a mixed-effects model, with time and group as fixed effects and individual mouse as a random effect. Statistical analysis was performed using GraphPad Prism. Differences were considered statistically significant at P value <0.05.

### Author contributions

T.Y. and K.H. contributed equally as co-first authors. The authorship order was determined because T.Y. assumed primary responsibility for writing the manuscript.

T.Y., K.H., J.T.W., and R.S.A. conceived the overall study. T.Y., K.H., J.B.L., J.L., T.J.L., J.C., A.S., and J.T.W. performed the experiments and analyzed the data. S.Y., R.T., B.S.S., D.D., K.K., and C.W.P. contributed methodology and resources. M.Y., J.T.W., and R.S.A. supervised the project and provided oversight. T.Y. and K.H. prepared all figures. T.Y., K.H., J.T.W., and R.S.A. wrote the manuscript with input from all authors. All authors reviewed the manuscript and approved the final version.

## Acknowledgements

We thank Dr. Lynn M. Hassman and Dr. Jonathan Kipnis for helpful discussions.

Experimental support was partially provided by the Bursky Center for Human Immunology and Immunotherapy Programs at Washington University, Immunomonitoring Laboratory. We also thank Washington University’s Siteman Flow Cytometry Core for help with flow sorting, Genome Technology Access Center for help with scRNA-seq studies, the Morphology & Imaging Core for assistance with paraffin sectioning and staining (Washington University Department of Ophthalmology & Visual Sciences).

Some illustrations were created using BioRender.com.

## Data Availability Statement

The scRNA-seq dataset generated by this study has been deposited in the Gene Expression Omnibus (GEO) under the accession number GSE296278 (Lin et al., 2025).

