## Supplemental data for "Macrophage immunosenescence prolongs intraocular inflammation in aged mice via impaired induction of regulatory T cells"

| Antibodies | Source | Identifier |
|---|---|---|
| AF488 anti-mouse CD25 Antibody (Clone PC61.5) | Thermo | Cat# 53-0251-80 |
| AF488 anti-mouse CD4 Antibody (Clone GK1.5) | BioLegend | Cat# 100425 |
| AF647 anti-mouse Ly6G Antibody (Clone 1A8) | BioLegend | Cat# 127609 |
| AF700 anti-mouse B220 Antibody (Clone RA3-6B2) | BioLegend | Cat# 103231 |
| APC anti-mouse TCRβ Antibody (Clone H57-597) | BioLegend | Cat# 109211 |
| APC/Cy7 anti-mouse CD11b Antibody (Clone M1/70) | BioLegend | Cat# 101225 |
| BV510 anti-mouse CD62L Antibody (Clone MEL-14) | BioLegend | Cat# 104441 |
| BV605 anti-mouse Ly6C Antibody (Clone HK1.4) | BioLegend | Cat# 128035 |
| Pacific Blue anti-mouse Helios Antibody (Clone 22F6) | BioLegend | Cat# 137210 |
| PE anti-mouse CD45 Antibody (Clone 30-F11) | BioLegend | Cat# 103105 |
| PE anti-mouse FOXP3 Antibody (Clone QA20A67) | BioLegend | Cat# 118903 |
| PE/Cy7 anti-mouse CD11c Antibody (Clone N418) | BioLegend | Cat# 117317 |
| PerCP/Cy5.5 anti-mouse CD64 Antibody (Clone X54-5/1.1) | BioLegend | Cat# 139307 |
| BUV395 anti-mouse NK1.1 Antibody (Clone PK136) | BD | Cat# 564144 |
| BV421 anti-mouse MHC II Antibody (Clone M5/114.15.2) | BD | Cat# 562564 |
| BV750 anti-mouse CD44 Antibody (Clone IM7) | BD | Cat# 747255 |
| BV786 anti-mouse CD69 Antibody (Clone H1.2F3) | BD | Cat# 564683 |
| PE/CF594 anti-mouse CD8a Antibody (Clone 53-6.7) | BD | Cat# 562283 |
| Purified Rat Anti-Mouse CD16/CD32 (Mouse BD Fc Block™) | BD | Cat# 553142 |

**Supplementary Table S1. Antibodies used in flow cytometry**

**2 weeks**

| | | | % in total cell counts | | % in CD45⁺ cells | |
|---|---|---|---|---|---|---|
| | Young | Old | Young | Old | Young | Old |
| Amacrine | 20 | 34 | 0.27% | 0.44% | | |
| Astrocytes | 39 | 99 | 0.52% | 1.27% | | |
| B | 37 | 3 | 0.49% | 0.04% | 0.67% | 0.11% |
| Bipolar | 115 | 399 | 1.53% | 5.12% | | |
| Cones | 133 | 510 | 1.77% | 6.55% | | |
| DC | 300 | 64 | 4.00% | 0.82% | 5.44% | 2.27% |
| Endothelial | 94 | 711 | 1.25% | 9.13% | | |
| Lymphocytes | 146 | 8 | 1.94% | 0.10% | 2.65% | 0.28% |
| **Mac/Mono** | **2106** | **706** | **28.05%** | **9.07%** | **38.18%** | **25.06%** |
| mature DC | 414 | 20 | 5.51% | 0.26% | 7.51% | 0.71% |
| Microglia | 635 | 1664 | 8.46% | 21.37% | 11.51% | 59.07% |
| Muller Glia | 438 | 1879 | 5.83% | 24.13% | | |
| Neutrophils | 471 | 72 | 6.27% | 0.92% | 8.54% | 2.56% |
| NK | 201 | 14 | 2.68% | 0.18% | 3.64% | 0.50% |
| Pericytes | 47 | 170 | 0.63% | 2.18% | | |
| Proliferating | 209 | 191 | 2.78% | 2.45% | 3.79% | 6.78% |
| Rods | 1097 | 977 | 14.61% | 12.55% | | |
| RPE | 8 | 191 | 0.11% | 2.45% | | |
| **T_CD4** | **901** | **53** | **12.00%** | **0.68%** | **16.33%** | **1.88%** |
| T_CD8 | 96 | 22 | 1.28% | 0.28% | 1.74% | 0.78% |
| Total | 7507 | 7787 | 100.00% | 100.00% | | |
| Total CD45⁺ cells | 5516 | 2817 | | | 100.00% | 100.00% |

**5 weeks**

| | | | % in total cell counts | | % in CD45⁺ cells | |
|---|---|---|---|---|---|---|
| | Young | Old | Young | Old | Young | Old |
| Amacrine | 59 | 50 | 0.74% | 0.61% | | |
| Astrocytes | 107 | 76 | 1.35% | 0.92% | | |
| B | 51 | 572 | 0.64% | 6.96% | 1.34% | 10.70% |
| Bipolar | 355 | 163 | 4.47% | 1.98% | | |
| Cones | 366 | 225 | 4.61% | 2.74% | | |
| DC | 484 | 558 | 6.09% | 6.79% | 12.68% | 10.44% |
| Endothelial | 193 | 222 | 2.43% | 2.70% | | |
| **Mac/Mono** | **878** | **1205** | **11.05%** | **14.65%** | **23.00%** | **22.54%** |
| mature DC | 238 | 462 | 3.00% | 5.62% | 6.23% | 8.64% |
| Microglia | 972 | 635 | 12.24% | 7.72% | 25.46% | 11.88% |
| Muller Glia | 1399 | 796 | 17.61% | 9.68% | | |
| Neutrophils | 72 | 66 | 0.91% | 0.80% | 1.89% | 1.23% |
| NK | 154 | 131 | 1.94% | 1.59% | 4.03% | 2.45% |
| Pericytes | 58 | 64 | 0.73% | 0.78% | | |
| Photoreceptors | 1066 | 942 | 13.42% | 11.46% | | |
| Proliferating | 212 | 198 | 2.67% | 2.41% | 5.55% | 3.70% |
| Rods | 441 | 297 | 5.55% | 3.61% | | |
| RPE | 66 | 22 | 0.83% | 0.27% | | |
| **T_CD4** | **504** | **908** | **6.35%** | **11.04%** | **13.20%** | **16.99%** |
| T_CD8 | 253 | 610 | 3.19% | 7.42% | 6.63% | 11.41% |
| Unknown | 15 | 21 | 0.19% | 0.26% | | |
| Total | 7943 | 8223 | 100.00% | 100.00% | | |
| Total CD45⁺ cells | 3818 | 5345 | | | 100.00% | 100.00% |

**Supplementary Table S2. Proportion of Immune cells identified by scRNA-seq**

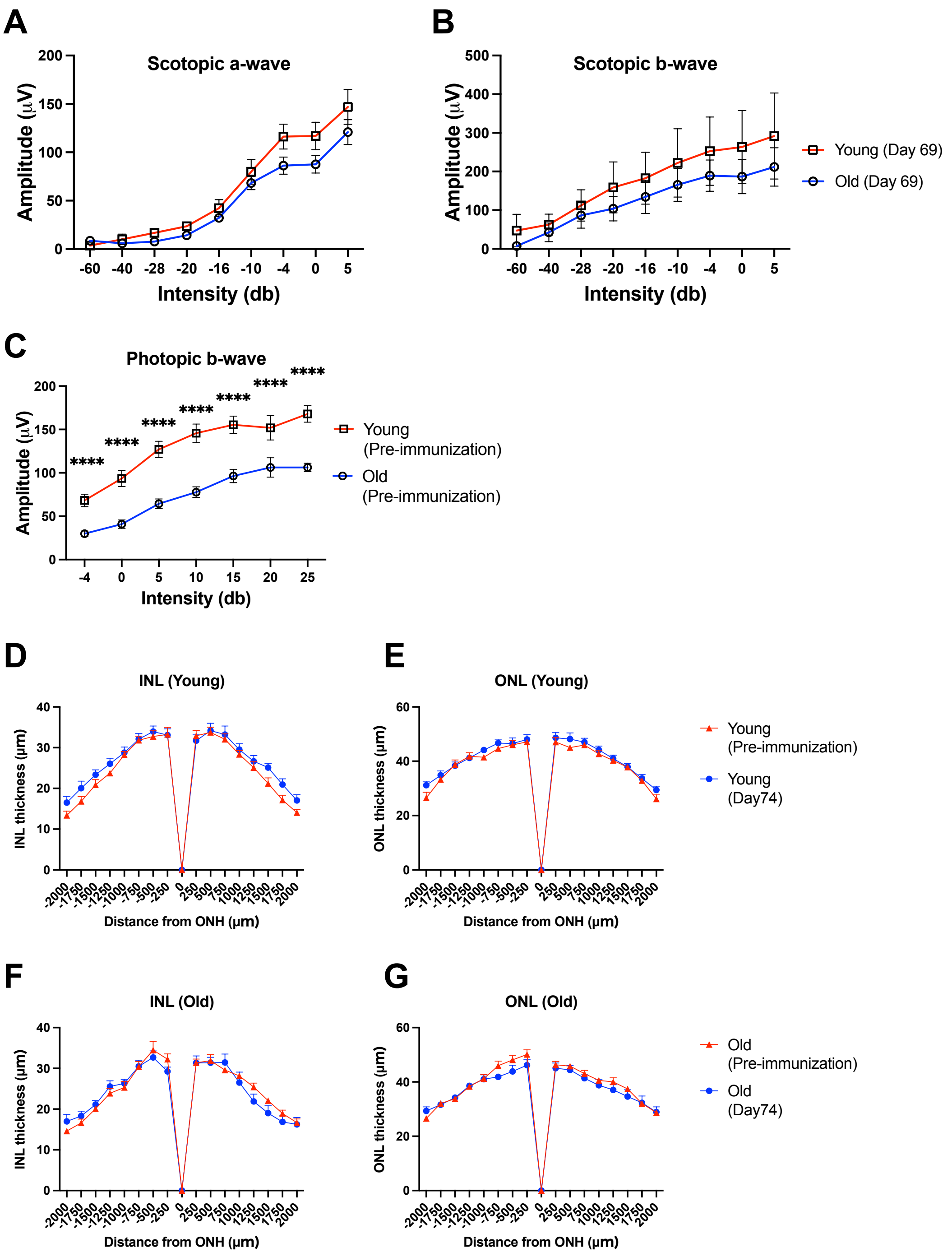


**Supplementary Figure S1. Preserved retinal thickness despite impaired visual function in the chronic phase of EAU**
**(A and B)** Full-field electroretinography assessing rod photoreceptor function (A, scotopic a-wave), rod-driven inner retina activity (B, scotopic b-wave), measured at day 69 (n = 8–12). Two-way ANOVA revealed a significant effect of age (column factor) on scotopic b-wave amplitudes ($P < 0.05$). **(C)** Full-field electroretinography of photopic b-wave measured before EAU induction (n = 8–12). **(D–G)** Retinal thickness of the inner nuclear layer (INL, D and F), representing intermediate neurons, and the outer nuclear layer (ONL, E and G), representing photoreceptor cell bodies, as measured by optical coherence tomography. No significant change in INL or ONL thickness was observed before versus after inflammation in either young or old mice. ****: $P < 0.0001$. Two-way ANOVA with Bonferroni's multiple comparisons test (A–G). All results are expressed as mean ± SEM.

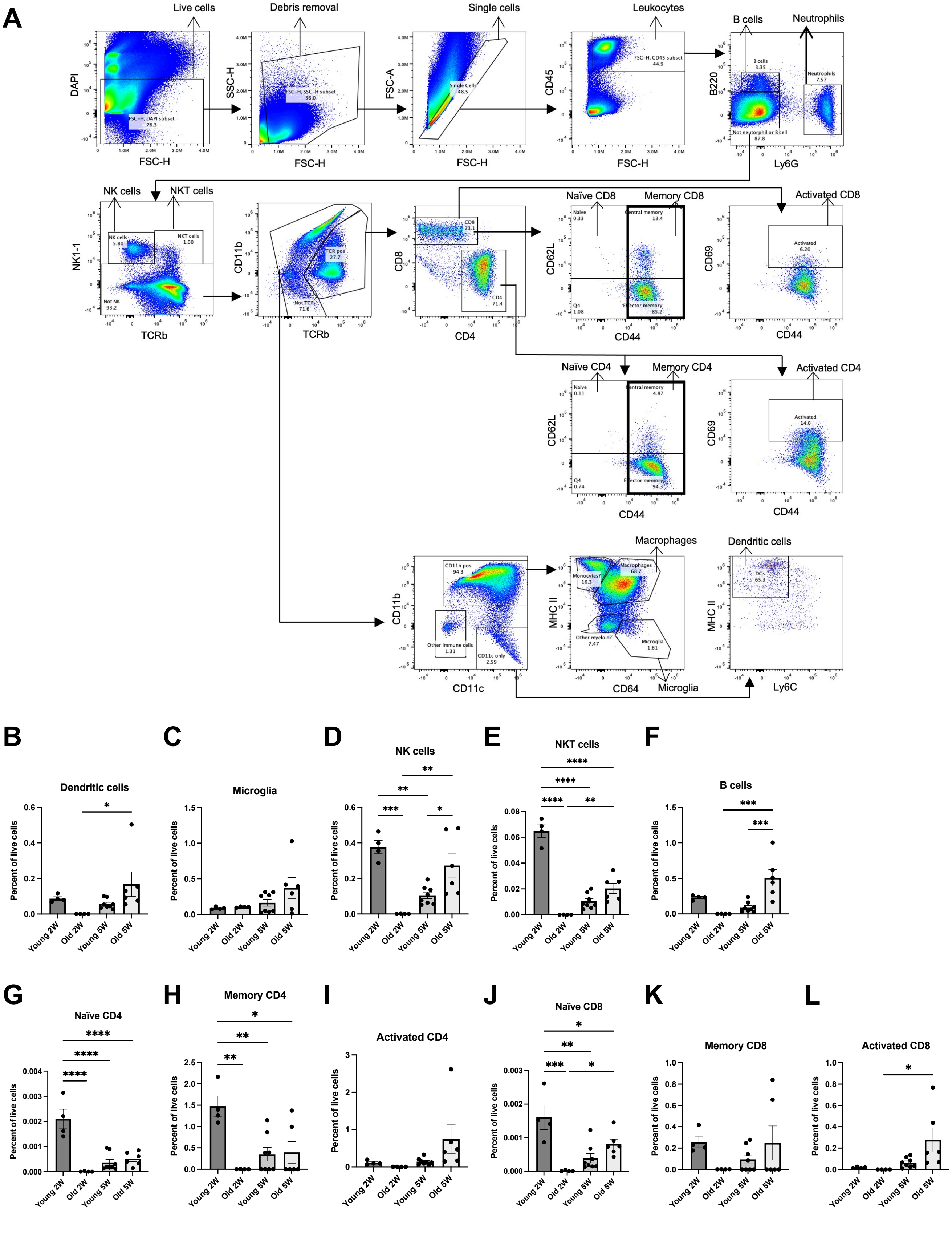


**Supplementary Figure S2. Gating strategy for high-dimensional flow cytometry and immune cell subset analysis**
**(A)** Gating strategy for high-dimensional flow cytometry. **(B–L)** Percentages of immune cell subsets (B–F) and T cell functional subsets (G–L) among total live cells in EAU retinas from young and old mice (Young 2W, n = 4; Old 2W, n = 4; Young 5W, n = 8; Old 5W, n = 6). *: $P < 0.05$; **: $P < 0.01$; ***: $P < 0.001$; ****: $P < 0.0001$. One-way ANOVA with Tukey's post hoc test (B–L). All results are expressed as mean ± SEM.

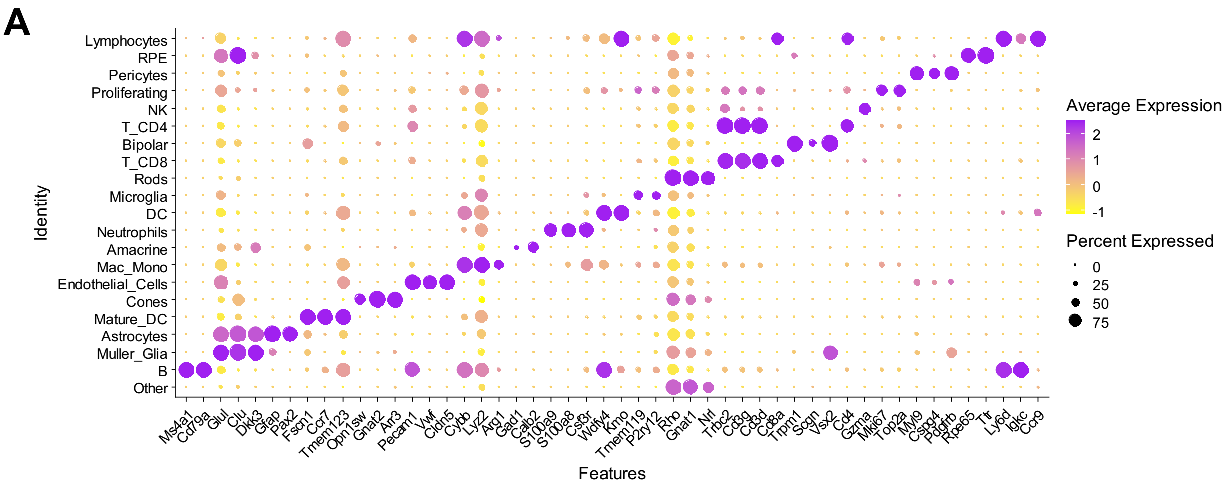


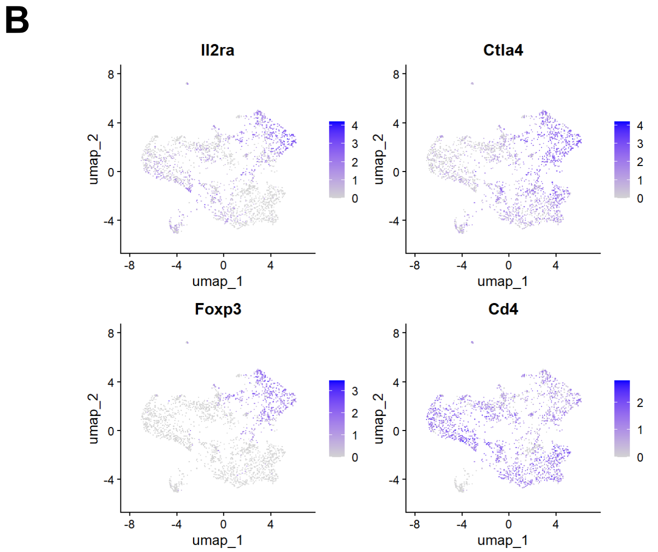


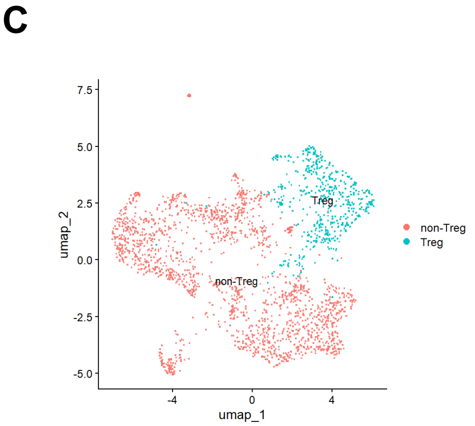


| | 2W_young | 2W_old | 5W_young | 5W_old |
|---|---|---|---|---|
| non-Treg | 755 | 47 | 347 | 682 |
| Treg | 146 | 6 | 157 | 226 |
| %Treg in total CD4+ | 16.20% | 11.32% | 31.15% | 24.89% |

**Supplementary Figure S3. Cell annotation and Treg re-clustering analysis in scRNA-seq dataset**
**(A)** Dot plot displaying the expression of canonical marker genes used for cell annotation of major immune cell populations and retinal parenchymal cell populations. **(B and C)** Treg re-clustering analysis in the T_CD4 population. UMAP plot showing the expression pattern of canonical Treg signature genes (B) and re-clustered Treg subset with the numbers of cells derived from each group (C).

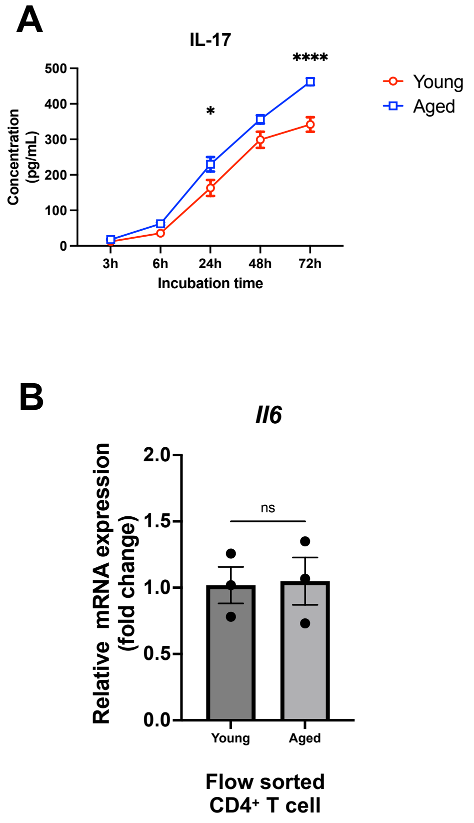


**Supplementary Figure S4. Cytokine profiling in co-culture supernatants and gene expression analysis in sorted $CD4^+$ T Cells** **(A)** IL-17 concentration in co-culture supernatants (n = 3–4). **(B)** Gene expression of IL-6 assessed in sorted $CD4^+$ T cells after 72 h of co-culture (n = 3). *: $P < 0.05$; ****: $P < 0.0001$; ns: not significant. Two-way ANOVA with Bonferroni's post hoc test (A). Unpaired Student's t test (B). All results are expressed as mean ± SEM.

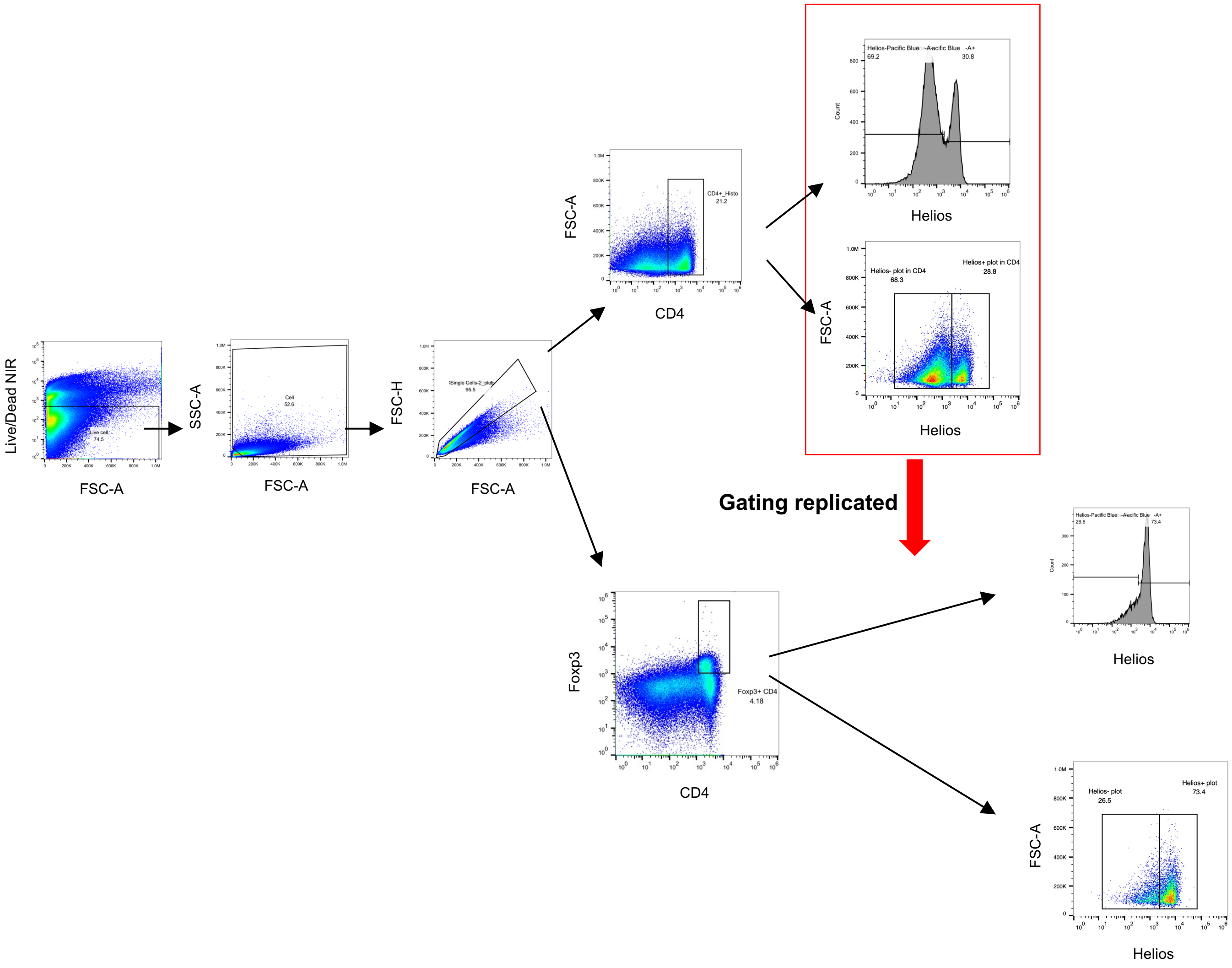


**Supplementary Figure S5. Gating strategy for Treg profiling in cervical lymph nodes**
Helios gating within Foxp3+ Tregs was adjusted based on the gating profile of total CD4+ T cells.
